# Detection and characterization of Langya virus in *Crocidura lasiura*, Republic of Korea

**DOI:** 10.1101/2024.12.02.626520

**Authors:** Augustine Natasha, Sarah E. Pye, Kyungmin Park, Shivani Rajoriya, Intae Yang, Jieun Park, Haryo Seno Pangestu, Jongwoo Kim, Yeonsu Oh, Carolina B. López, Jin-Won Song, Won-Keun Kim

## Abstract

This study reports the identification of Langya virus Korea (LayV KOR) during the surveillance of shrews in the Republic of Korea. LayV KOR represents the first identification of LayV outside of China, exhibiting approximately 80% and 95.5% homologies at nucleotide and amino acid levels.

## Introduction

The *Henipavirus* is a genus that includes the highly virulent and re-emergent Nipah and Hendra viruses whose mortality rates can exceed 70% in humans (1). Related to these viruses, Langya virus (LayV) is a zoonotic *Henipavirus* first identified in 2018 from patients with fever in China (2). The shrew species *Crocidura lasiura* and *C. shantungensis* were found to be natural hosts of LayV. Our group previously documented the presence of the Henipa-like viruses, Gamak virus (GAKV) and Daeryeong virus (DARV), in shrews in the Republic of Korea (ROK) (3). However, LayV has not been reported outside China.

In this study, we delineated for the first time the complete genome sequence of LayV identified in *C. lasiura* during surveillance monitoring in the ROK, using a combination of metagenomic and amplicon-based sequencing. Through retrospective analysis of prior sequencing data, we identified Langya virus sequences from Korean shrew samples as early as 2017. Analysis of these genome sequences revealed the presence of a previously unreported hypothetical protein domain between the matrix and fusion proteins. Phylogenetic analysis demonstrated a close evolutionary relationship between LayVs from the ROK and China. This is the first report of LayV outside China, highlighting the cross-border circulation of emerging shrew-borne henipaviruses.

### The Study

In 2023, we collected kidney tissues from 24 *C. lasiura* and screened them for paramyxoviruses using the reverse transcription PCR method described previously (4, 5). The animal handling was performed according to the Korea University Institutional Animal Care and Use Committee (#2022□34). The specimens were collected from the Gyeonggi, Gangwon, Chungcheongnam, and Jeollanam Provinces (Supplementary Figure 1). Detailed sample characteristics are listed in Table 1. The positivity rate for paramyxovirus was 62.5%, with a higher rate in males than in females (76.9% vs. 45.5%). The captured animals exhibited a high positivity rate in the adult group (75%).

**Table 1.**
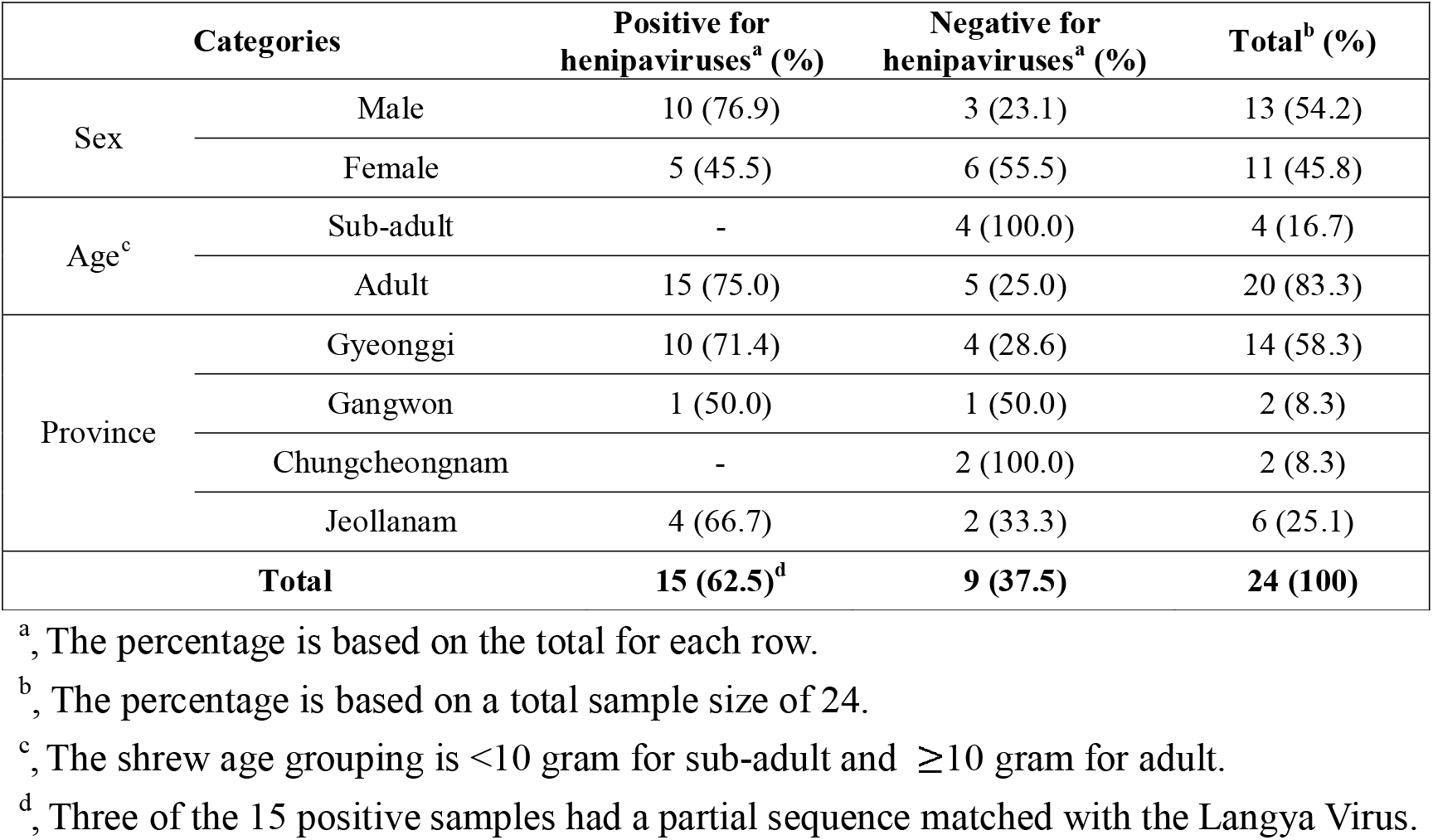
Prevalence of henipaviruses in *Crocidura lasiura* samples collected in the ROK in 2023.

Geographically, we obtained more samples from the northern than the southern areas, with the highest positivity rate (71.4%) in Gyeonggi Province. We aligned the partial sequences using MUSCLE 5, followed by phylogenetic analysis using IQTREE and the Interactive Tree of Life (ITOL) (6–8). Notably, 3 of 15 (20%) partial sequences from the ROK shared a common ancestor with LayV from China (Supplementary Figure 2).

While the first report of LayV didn’t emerge until 2018, we retrospectively analyzed the metagenomic data from our previous study in 2017, which led to the discovery of GAKV in *C. lasiura* (3). We focused on Cl17-6 and Cl17-9, whose partial sequences clustered phylogenetically independent of GAKV. De novo assembly was performed using the QIAGEN CLC Genomic Workbench (https://digitalinsights.qiagen.com), resulting in long contigs that exhibited 79.7% and 79.8% nucleotide similarities to LayV China. These sequences were subsequently designated as LayV KOR Cl17-6 and Cl17-9. Annotation of the open reading frames (ORFs) from 3□ to 5□ (Figure 1) showed the genomic organization as nucleocapsid (N), phosphoprotein (P), matrix protein (M), hypothetical protein (h), fusion protein (F), glycoprotein (G), and large protein (L). The 3□ leader and 5□ trailer sequences were determined using Rapid Amplification of cDNA Ends PCR with the SMARTer RACE 5□/3□ (TaKaRa Bio Inc., Japan). The complete genome sequence of LayV KOR was 18,420 nucleotides (nt) in length, with the leader sequence ACCAAA at the 3□ end and the trailer sequence TGTGGT at the 5□ end. Based on this sequence, we designed primer sets for amplicon-based sequencing using PrimalScheme (9). cDNA from the kidney tissue of *C. lasiura* captured in 2023 was synthesized and used as a template for tiled PCR following the ARTIC Network protocol (https://artic.network/2-protocols.html). Amplicon-based sequencing was performed on a MinION platform with R10 flow cell chemistry and the Native Barcoding Kit 24 V14 (Oxford Nanopore Technologies, UK). Raw reads were demultiplexed, trimmed, and mapped to the LayV KOR Cl17-6 as a reference sequence using the QIAGEN CLC Genomic Workbench toolkit. The consensus sequence was annotated based on the henipaviruses from the NCBI database. Three complete sequences from 2023 samples were designated as LayV KOR Cl23-2, Cl23-18, and Cl23-19, respectively.

**Figure 1.**
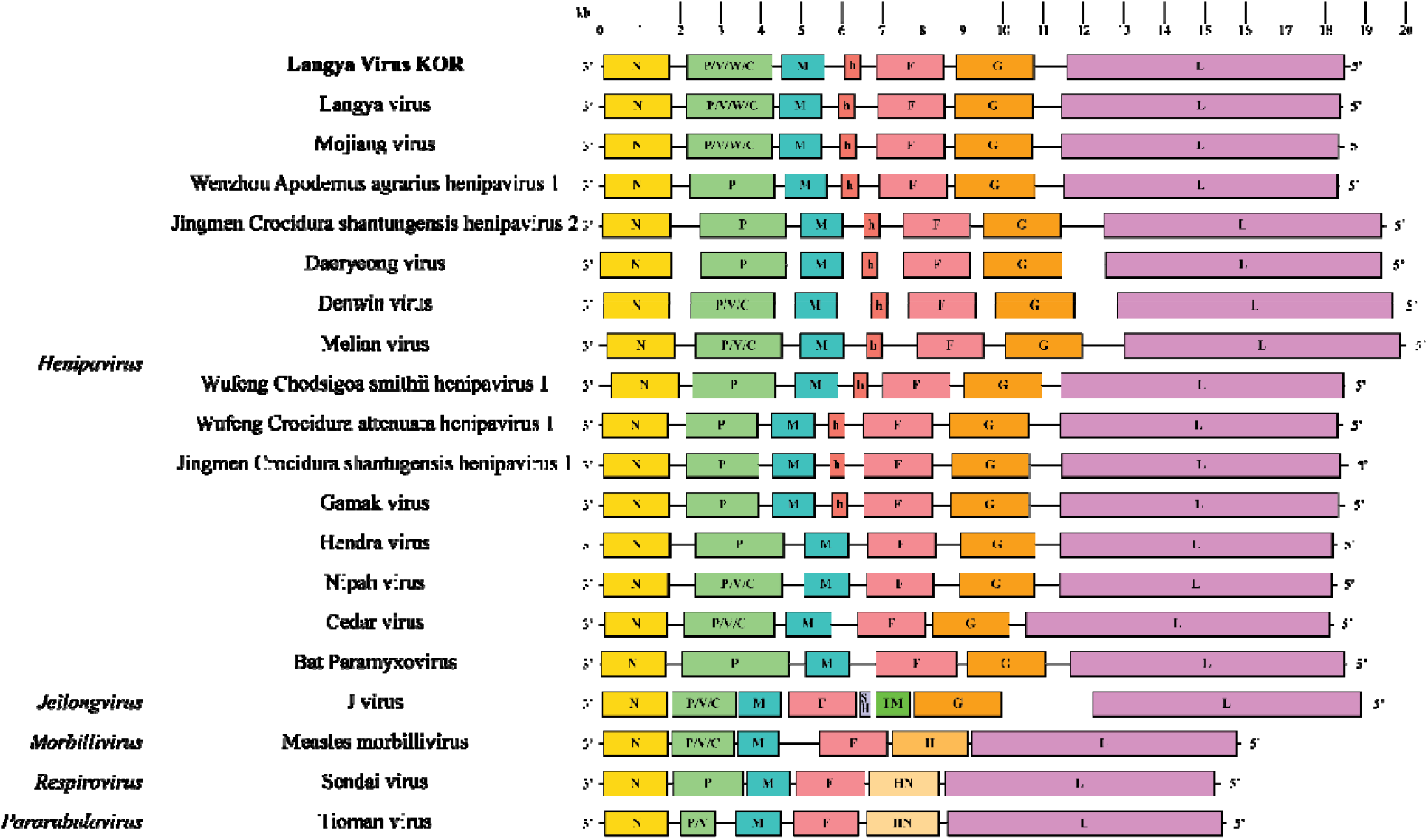
The genome organization of Langya virus Korea (LayV KOR) compared with the LayV from China and other reported henipaviruses. Abbreviations: N, nucleocapsid protein; P, phosphoprotein; M, matrix protein; h, hypothetical protein; F, fusion protein; SH, small hydrophobic protein; TM, transmembrane protein; G, glycoprotein; H, hemagglutinin protein; HN, hemagglutinin-neuraminidase protein; L, large protein.

We obtained complete genomes that fulfilled the rule of six, a requisite for productive paramyxovirus genomes. The full genome sequence length was 18 nt longer than that of LayV from China, with variations in the non-coding area. Our analysis identified a 366 nt ORF, designated as a “hypothetical protein” (h) between the M and F genes. This ORF has also been reported in shrew-borne henipaviruses but is absent in bat-borne henipaviruses (Figure 1). Notably, the canonical gene start signal for the hypothetical protein was located after the M gene stop signal and intergenic region (IGR), and a non-canonical stop-IGR-start signal was identified between the h- and F-coding sequences. The biological significance of this hypothetical protein remains to be investigated.

Phylogenetic relationship of the five complete LayV KOR genome sequences shared a common ancestor but exhibited a distinct genetic lineage with the LayV China (Figure 2). Similarity analysis of the coding sequences (CDSs) and their translated protein sequences compared to LayV from China demonstrated average similarities of 83.0% and 95.5%, respectively. The nucleotide similarity dropped to an average of 53.3% and 62.2%, compared with the CDS of GAKV and DARV. At the amino acid level, the similarity of these shrew-borne henipaviruses to LayV KOR was even lower, ranging from 47.1% for GAKV to 60.4% for DARV (Figure 3). The phylogenetic inference and high sequence similarities of LayV between ROK and China indicate a closer evolutionary relationship compared to other shrew-borne henipaviruses found in the ROK.

**Figure 2.**
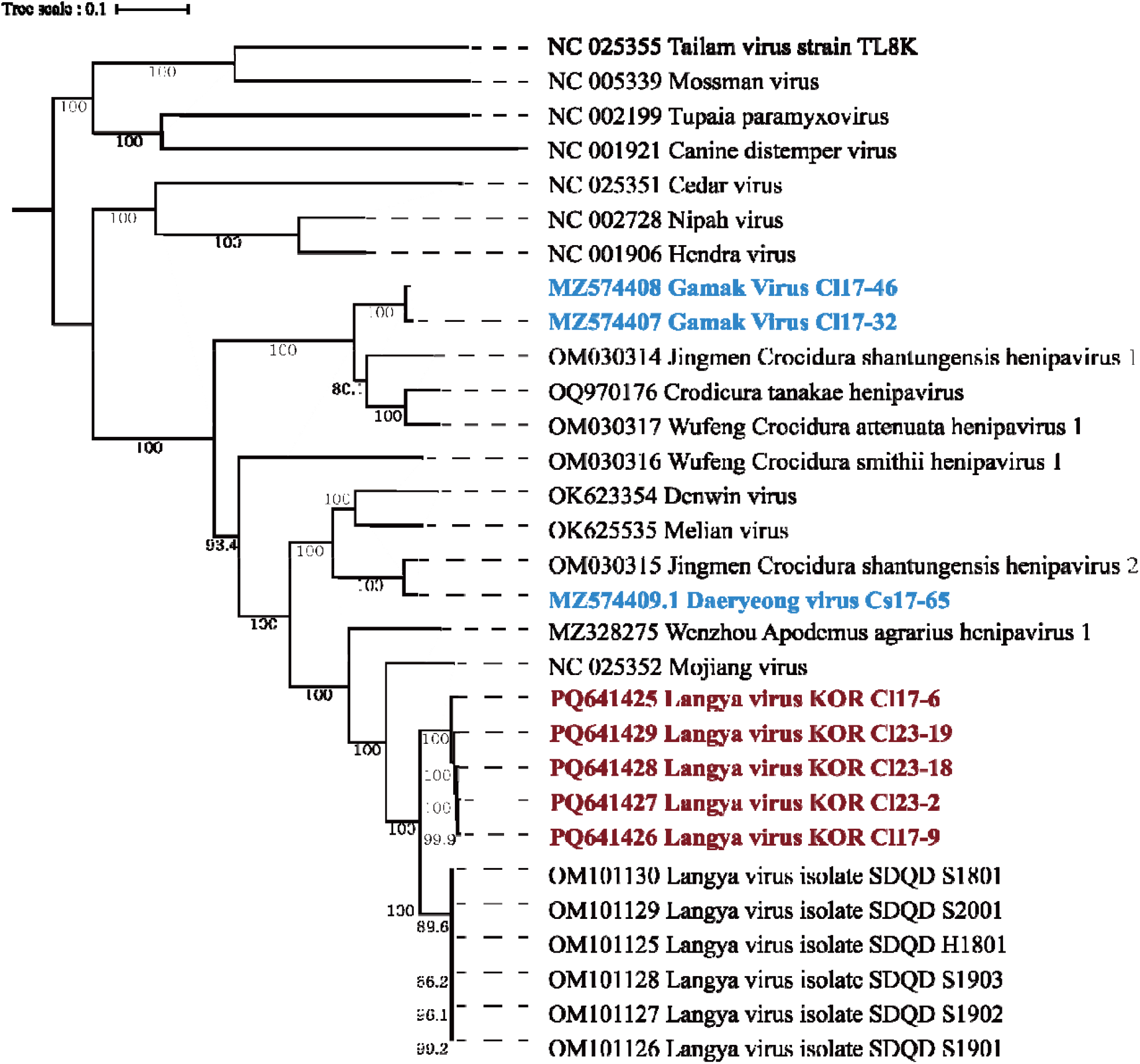
Phylogenetic tree of the Langya virus Korea (LayV KOR) complete sequences aligned with other paramyxoviruses. The tree was constructed using maximum likelihood analysis via IQTREE web server with the GTR+F+I+G4 model selected based on BIC and 1000 bootstrapping. The burgundy-colored labels represent the LayV KOR reported in this study, while the blue-colored labels represent the shrew-borne henipaviruses found in the Republic of Korea from a previous study.

**Figure 3.**
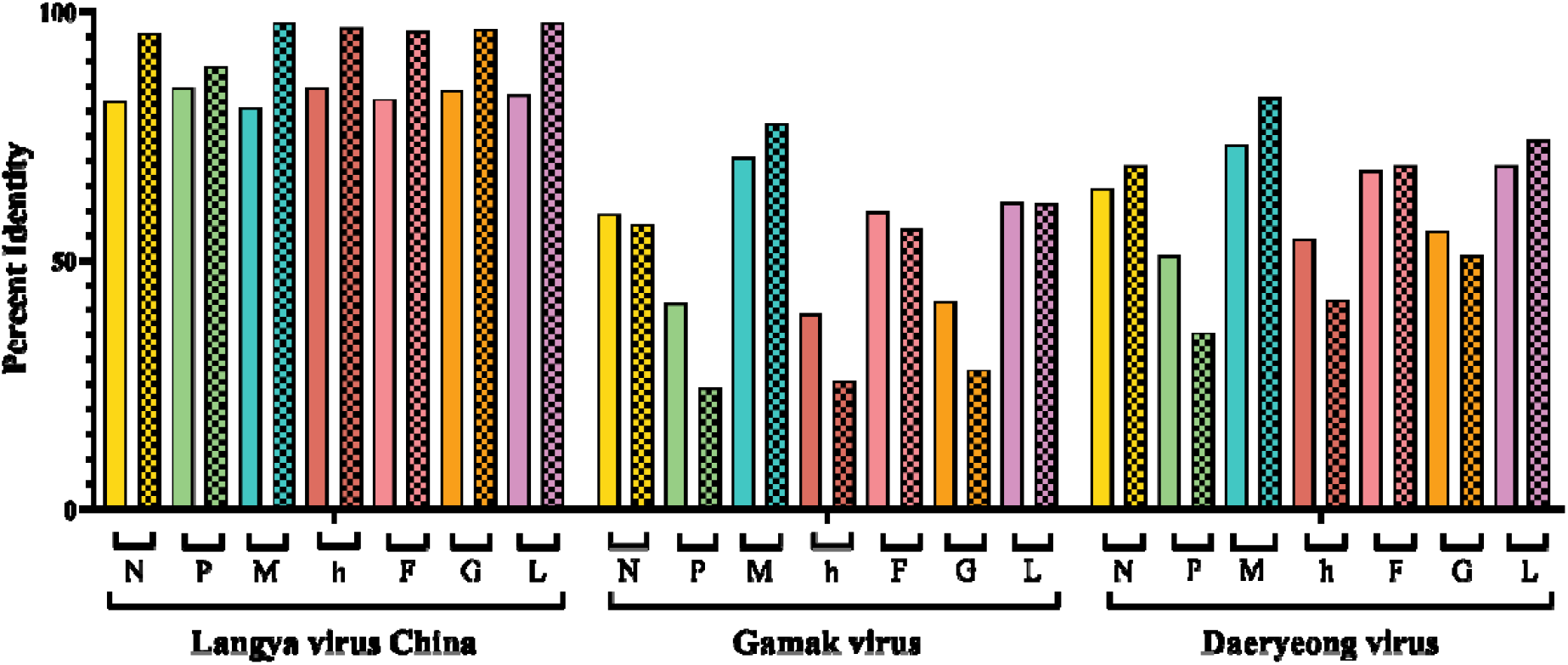
Percentage similarity of the genomic and amino acid sequences of the Langya virus Korea (LayV KOR) coding sequence against LayV China, Gamak virus (GAKV), and Daeryeong virus (DARV). The bar with a solid color represents the nucleotide sequence of the gene, whereas the bar with a checkered pattern represents the translated gene.

LayV infection was first identified in clinical samples collected in 2018, whereas our group observed complete sequences in shrew samples collected in 2017 and 2023. To date, no human-to-human transmission or deaths associated with LayV infection have been reported. The current number of clinical cases remains low, limiting the ability to assess the risk to vulnerable groups.

LayV infection lacks pathognomonic signs, with fever being its primary symptom. The index case of LayV was detected through active surveillance using metagenomic sequencing in patients with fever and a history of animal exposure. This led to a hypothesis that LayV KOR infections in the Korean population might have been underdiagnosed since the sentinel monitoring is infrequent. Initial LayV cases were predominantly found in individuals working as farmers (85%), suggesting that outdoor activities might serve as a potential route for human exposure to shrews. In humans, LayV was detected using throat swabs, indicating viral shedding in the respiratory tract. Similar to other respiratory viruses of zoonotic origin, including betacoronaviruses and influenza viruses, rapid transmission of LayV may become unavoidable once the virus adapts to infect humans (10, 11).

This study has several limitations: the current absence of viral isolates, the limited geographic surveillance and the unexplored potential for human exposure. The tropism of LayV KOR may hinder effective virus isolation using common cell lines requiring the need for additional tool development.

## Conclusion

Our findings provide evidence for the circulation of LayV in *C. lasiura* in the ROK, which closely related to a zoonotic pathogen previously identified exclusively in China. This report raises awareness and caution among physicians regarding clinical cases of emerging shrew-borne henipaviruses. Future studies should assess evidence of LayV KOR exposure in humans by developing antibody screening for active serosurveillance in the ROK.

## Acknowledgments

We thank Dr. David Wang (Washington University, St. Louis, USA) for the critical discussions. We thank Nayeon Jang (Hallym University) for supporting this study.

## Data Availability

All the sequences are available in the NCBI GenBank database with accession number PQ641425-PQ641429.

## Funding

This study was supported by the Basic Research Program through the National Research Foundation of Korea (NRF) by the Ministry of Education (NRF-2021R1I1A2049607) and Government-wide R&D to Advance Infectious Disease Prevention and Control, Republic of Korea (RS-2023 - KH140418). This work was supported by the NRF Korea grant funded by the Korean government (MSIT) (2023R1A2C2006105). This research was also supported by a Novo Nordisk Foundation PAD award (NF22SA0082041) and BJC investigator funds. This study was partly supported by the NIH grant (U01 AI151810).

## Disclosure

The authors declare no competing financial interests.

---

**Supplementary figure 1.**
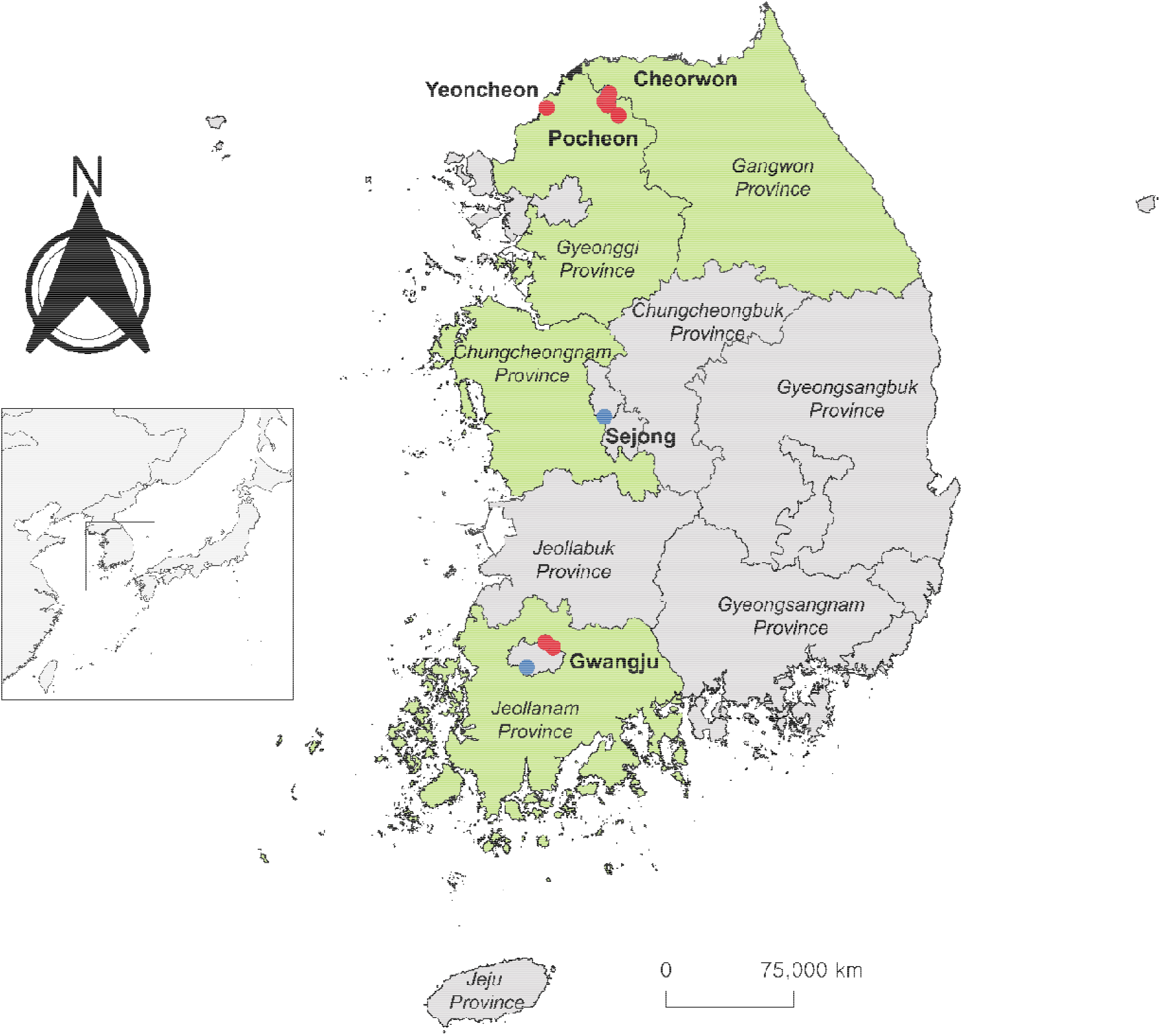
The map shows the trapping locations for collecting *C. lasiura* in 2023. The green-colored provinces indicate areas covered in the 2023 shrew surveillance project. The red-colored dots represent areas with paramyxovirus-positive samples, while the blue-colored dots represent areas with paramyxovirus-negative samples.

**Supplementary figure 2.**
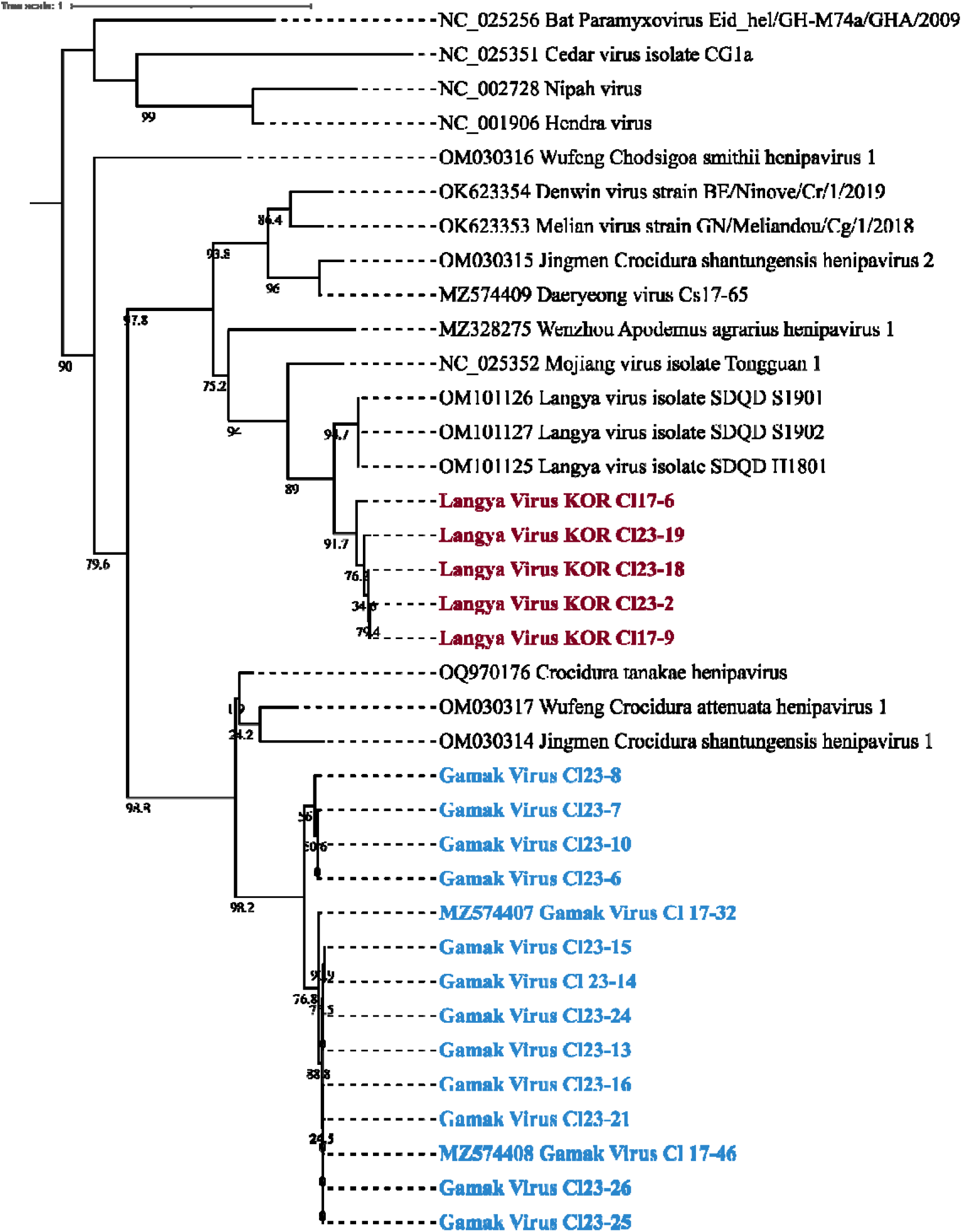
The phylogenetic tree of the collected partial sequences from paramyxovirus PCR screening. The tree was constructed using maximum likelihood analysis by IQTREE web server, with GTR+F+I+G4 model chosen according to BIC and 1000 bootstrapping. Burgundy-colored labels represent samples with partial sequences identified as Langya Virus KOR, while the blue colored labels represent partial sequences identified as Gamak Virus.

## Notes

### Competing Interest Statement

The authors have declared no competing interest.

### Summary of Updates

We revise the funding identification code which supporting this study.

